# Bayesian model comparison for rare variant association studies

**DOI:** 10.1101/257162

**Authors:** Guhan Ram Venkataraman, Christopher DeBoever, Yosuke Tanigawa, Matthew Aguirre, Alexander G. Ioannidis, Hakhamanesh Mostafavi, Chris C. A. Spencer, Timothy Poterba, Carlos D. Bustamante, Mark J. Daly, Matti Pirinen, Manuel A. Rivas

## Abstract

Whole genome sequencing studies applied to large populations or biobanks with extensive phenotyping raise new analytic challenges. The need to consider many variants at a locus or group of genes simultaneously and the potential to study many correlated phenotypes with shared genetic architecture provide opportunities for discovery and inference that are not addressed by the traditional one variant, one phenotype association study. Here, we introduce a Bayesian model comparison approach that we refer to as MRP (Multiple Rare-variants and Phenotypes) for rare-variant association studies that considers correlation, scale, and direction of genetic effects across a group of genetic variants, phenotypes, and studies. The approach requires only summary statistic data. To demonstrate the efficacy of MRP, we apply our method to exome sequencing data (N = 184,698) across 2,019 traits from the UK Biobank, aggregating signals in genes. MRP demonstrates an ability to recover previously-verified signals such as associations between *PCSK9* and LDL cholesterol levels. We additionally find MRP effective in conducting meta-analyses in exome data. Notable non-biomarker findings include associations between *MC1R* and red hair color and skin color, *IL17RA* and monocyte count, *IQGAP2* and mean platelet volume, and *JAK2* and platelet count and crit (mass). Finally, we apply MRP in a multi-phenotype setting; after clustering the 35 biomarker phenotypes based on genetic correlation estimates into four clusters, we find that joint analysis of these phenotypes results in substantial power gains for gene-trait associations, such as in *TNFRSF13B* in one of the clusters containing diabetes and lipid-related traits. Overall, we show that the MRP model comparison approach is able to improve upon useful features from widely-used meta-analysis approaches for rare variant association analyses and prioritize protective modifiers of disease risk.

## Introduction

Sequencing technologies are quickly transforming human genetic studies of complex traits. It is increasingly possible to obtain whole genome sequence data on thousands of samples at manageable costs. As a result, the genome-wide study of rare variants (minor allele frequency [MAF] < 1%) and their contribution to disease susceptibility and phenotype variation is now feasible.^1–4^

In genetic studies of diseases or continuous phenotypes, rare variants are hard to assess individually due to the limited number of observations of each rare variant. Hence, to boost the power to detect a signal, evidence is usually aggregated across variants in blocks. When designing an aggregation method, there are three questions that are usually considered. First, across which biological units should variants be combined (e.g. genes); second, which variants within those units should be included^5^; and third, which statistical model should be used?^6^ Given the widespread observations of shared genetic risk factors across distinct diseases, there is also considerable motivation to use gene discovery approaches that leverage the information from multiple phenotypes jointly. In other words, rather than only aggregating variants that may have effects on a single phenotype, we can also bring together sets of phenotypes for which a single variant or set of variants might have effects.

In this paper, we present a Bayesian **M**ultiple **R**are-variants and **P**henotypes (MRP) model comparison approach for identifying rare-variant associations as an alternative to current, widely-used univariate statistical tests. The MRP framework exploits correlation, scale, and/or direction of genetic effects in a broad range of rare-variant association study designs including case-control, multiple diseases and shared controls, a single continuous phenotype, multiple continuous phenotypes or a mixture of case-control and multiple continuous phenotypes (**Figure 1**). MRP makes use of Bayesian model comparison, whereby we compute a Bayes Factor (BF) defined as the ratio of the marginal likelihoods under two models: 1) a null model where all genetic effects are zero; and 2) an alternative model where factors like correlation, scale and direction of genetic effects are considered. For MRP, the BF represents the statistical evidence for a non-zero effect for a particular group of rare variants on the phenotype(s) of interest and can be used as an alternative to p-values from traditional significance testing.

**Figure 1.**
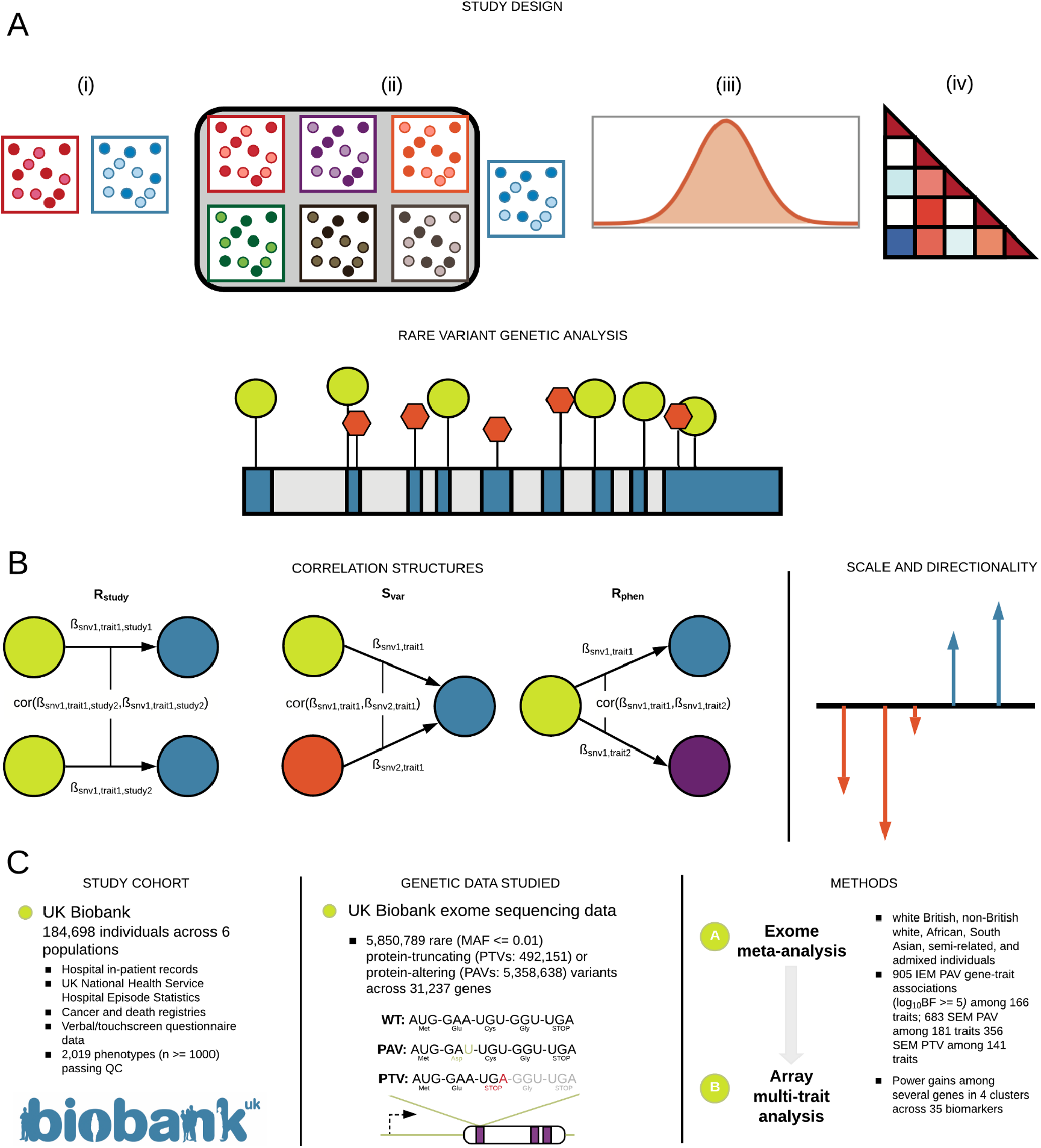
MRP study overview. **1A)** MRP is suitable for a broad range of rare variant association study designs, including, from left to right: **i)** case-control, **ii)** multiple diseases with shared controls, **iii)** single quantitative phenotype, and **iv)** mixtures of case-control and quantitative phenotypes. **1B)** Diagram of factors considered in rare variant association analysis including the correlation matrices: ***R***_***study***_ (expected correlation of genetic effects among a group of studies), ***S***_***var***_ (expected covariance of genetic effects among a group of variants, potentially accounting for annotation of variants), and ***R***_***phen***_ (expected correlation of genetic effects among a group of phenotypes). MRP can take into account both scale and direction of effects. **1C)** We focused on 184,698 individuals across 6 ancestry groups in the UK Biobank and analyzed 5,850,789 rare coding variants (492,151 PTVs, 5,358,638 PAVs) in the whole exome sequencing data via single-trait and multi-trait meta-analyses, with a specific focus on 35 biomarker traits.

While many large genetic consortia collect both raw genotype and phenotype data, in practice, sharing of individual genotype and phenotype data across groups is difficult to achieve. To address this, MRP can use summary statistics, such as estimates of effect size and corresponding standard errors from typical single-variant/single-phenotype linear or logistic regressions, as input. Furthermore, we use insights from Liu et al.^7^ and Cichonska et al.,^8^ which suggest the use of additional summary statistics like covariance estimates across variants and studies, respectively, for the lossless ability to detect gene-based association signals using summary statistics alone.

Aggregation techniques rely on variant annotations to assign variants to groups for analysis. MRP allows for the inclusion of priors on the scale of effect sizes that can be adjusted depending on what type of variants are included in the analysis. For instance, protein truncating variants (PTVs)^9,10^ are highly likely to be functional because they often disrupt the normal function of a gene. Additional deleteriousness metrics, such as MPC (which combines subgenic constraints with variant-level data for deleteriousness prediction)^11^ and pLI (derived from a comparison of the observed number of PTVs in a sample to the number expected in the absence of fitness effects, i.e. under neutrality, given an estimated mutation rate for the gene)^12^, can further attenuate or accentuate these granular signals. Furthermore, since PTVs typically abolish or severely alter gene function, there is particular interest in identifying protective PTV modifiers of human disease risk that may serve as targets for future therapeutics.^13–15^ We therefore demonstrate how the MRP model comparison approach can improve discovery of such protective signals by modeling the direction of genetic effects; this prioritizes variants or genes that are consistent with protecting against disease.

To evaluate the performance of MRP, we use simulations and compare it to other commonly used approaches. Some simple alternatives to MRP include univariate approaches for rare variant association studies including the sequence kernel association test (SKAT)^16^, and the burden test^6^, which are special cases of the MRP model comparison when we assign the prior correlation of genetic effects across different variants to be zero or one, respectively.

We apply MRP to summary statistics computed on a tranche of N = 184,698 exomes for thousands of traits in the UK Biobank for which we have exome data for N ≥ 1000 white British individuals, focusing on a meta-analysis context across six UK Biobank subpopulations as defined previously (**Methods**).^17^ We additionally apply multi-phenotype MRP on clusters of biomarker traits within a single-population context (white British individuals). These analyses show that MRP recovers results from single variant-single phenotype association analyses while increasing the power to detect new rare variant associations, including protective modifiers of disease risk.

## Methods

### Description of MRP

In this section, we provide an overview of the MRP model comparison approach (the **Supplementary Note** contains additional details). MRP models GWAS summary statistics as being distributed according to one of two models: the null model, where the effect sizes across all studies for a group of variants and a group of phenotypes is zero, and the alternative model, where effect sizes are distributed according to a multivariate normal distribution with a non-zero mean and/or covariance matrix. MRP compares the evidence between the alternative model and the null model using a Bayes Factor (BF) that is the ratio of the marginal likelihoods under the two models given the observed data.

To define the alternative model, we must specify the prior correlation structure, scale, and direction of the effect sizes. Let *N* be the number of individuals and *K* the number of phenotype measurements on each individual. Let *M* be the number of variants in a testing unit *G*, where *G* can be, for example, a gene, pathway, or a network. Let *S* be the number of studies from which data is obtained 一 this data may be in the form of **a)** raw genotypes and phenotypes, or **b)** summary statistics including linkage-disequilibrium coefficients, effect sizes, and corresponding standard errors. When considering multiple studies (*S* > 1), multiple rare variants (*M* > 1), and multiple phenotypes (*K* > 1), we define the prior correlation structure of the effect sizes as an *SMK* x *SMK* matrix, *U*. In practice, we define *U* as a Kronecker product, of three sub-matrices:

- an *S* x *S* matrix ***R***_***study***_ containing the correlations of genetic effects among studies that can model the level of heterogeneity in effect sizes between populations^18^;
- an *M* x *M* matrix ***S***_***var***_ containing the covariances of genetic effects among genetic variants, which may reflect, e.g., the assumption that all the PTVs in a gene may have the same biological consequence^9,10,19^ or prior information on scale of the effects obtained through integration of additional functional data ^5,20^; by assuming zero correlation of genetic effects, MRP becomes a dispersion test similar to C-alpha^21,22^ and SKAT^16^; and
- a *K* x *K* ***R***_***phen***_ matrix containing the correlations of genetic effects among phenotypes, which may be estimated from common variant data.^23–25^

The variance-covariance matrix of the effect size estimates may be obtained from readily available summary statistics such as in-study LD matrices, effect size estimates (or log odds ratios), and the standard errors of the effect size estimates (**Supplementary Note**).

MRP allows users to specify priors that reflect knowledge of the variants and phenotypes under study. For instance, we can define an independent effects model (IEM) where the effect sizes of different variants are not correlated at all. In this case, ***S***_***var***_ is the identity matrix, and MRP behaves similarly to dispersion tests like C-alpha^21,22^ and SKAT^16^. We can also define a similar effects model (SEM) by setting every value of ***R***_***var***_ to ∼ 1, where ***R***_***var***_ is the correlation matrix corresponding to covariance matrix ***S***_***var***._ This model assumes that all variants under consideration have similar effect sizes (with, possibly, differences in scale; like in the burden test). Such a model may be appropriate for PTVs, where each variant completely disrupts the function of the gene leading to a gene knockout. The prior on the scale of effect sizes can be used to denote which variants may have larger effect sizes. For instance, emerging empirical genetic studies have shown that within a gene PTVs may have stronger effects than missense variants.^26^ This can be reflected by adjusting the prior variances of effect sizes (σ) for different categories of variants (**Supplementary Note**).

Finally, we can utilize a prior on the expected location and direction of effects to specify alternative models where we seek to identify variants with protective effects against disease. By default, we have assumed that the prior mean, of genetic effects is zero, which makes it possible to analyze a large number of phenotypes without enumerating the prior mean across all phenotypes. To proactively identify genetic variants that are consistent with a protective profile for a disease, we can include a non-zero vector as a prior mean of genetic effects (**Supplementary Note**). For this, we can exploit information from Mendelian randomization studies of common variants, such as recent findings where rare protein-truncating loss-of-function variants in *PCSK9* were found to decrease LDL and triglyceride levels and decrease CAD risk^13,27,28^ to identify situations where such a prior is warranted.

Applying MRP to variants from a testing unit *G* yields a BF for that testing unit that describes the evidence that rare variants in that testing unit have a nonzero effect on the traits used in the model. We can turn this evidence into probability via Bayes rule. Namely, a multiplication of prior-odds of association by BF transforms the prior-odds to posterior-odds. For example, if our prior probability for one particular gene to be associated with a phenotype is 10^−4^, then an observed BF of 10^5^ means that our posterior probability of association between the gene and the phenotype is over 90%. Although we see advantages in adopting a Bayesian interpretation for MRP, our approach could also be used in a frequentist context by using BF as a test statistic to compute *p*-values (**Supplementary Note**).

### UK Biobank Data

#### Population definitions

**Table 1.** Number of individuals per population per genotyping platform (exome/array).

| Population | $N_{\text{exome}}$ | $N_{\text{array}}$ |
| --- | --- | --- |
| White British | 137,920 | 337,138 |
| Non-British White | 10,432 | 24,905 |
| African | 2,716 | 6,497 |
| South Asian | 3,569 | 7,885 |
| Semi-Related | 18,100 | 44,632 |
| Admixed | 11,961 | 28,551 |
| Total | 184,698 | 449,608 |

We used a combination of self-reported ancestry (UK Biobank field ID 21000) and principal component analysis to identify six subpopulations in the study: white British, African, South Asian, non-British white, semi-related, and an admixed population. To determine the first four populations, which contain samples not related closer than the 3rd degree, we first used the principal components of the genotyped variants from the UK Biobank and defined thresholds on principal component 1 and principal component 2 and further refined the population definition.^17^ Semi-related individuals were grouped as individuals whose genetic data (after passing UK Biobank QC filters; sufficiently low missingness rates; and genetically inferred sex matching reported sex), using a KING relationship table, were between conditional third and conditional second degrees of relatedness to samples in the first four groups. Admixed individuals were grouped as unrelated individuals who were flagged as “used_in_pca_calculation” by the UK Biobank and were not assigned to any of the other populations.

#### GWAS Summary Statistics

We performed genome-wide association analysis on 2,019 UK Biobank traits in the six population subgroups as defined above using PLINK v2.00a (20 October 2020). We used the --glm Firth fallback option in PLINK to apply an additive-effect model across all sites. Quantitative trait values were rank normalized using the --pheno-quantile-normalize flag. We used the following covariates in our analysis: age, sex, array type, and the first ten genetic principal components, where array type is a binary variable that represents whether an individual was genotyped with UK Biobank Axiom Array or UK BiLEVE Axiom Array. For variants that were specific to one array, we did not use array as a covariate.

For the admixed population, we conducted local ancestry-corrected GWAS. We first assembled a reference panel from 1,380 single-ancestry samples in the 1000 Genomes Project,^29^ the Human Genome Diversity Project,^30^ and the Simons Genome Diversity Project,^31^ choosing appropriate ancestry clusters by running ADMIXTURE^32^ with the unsupervised setting. Using cross-validation, eight well-supported ancestral population clusters were identified: African, African Hunter-Gatherer, East Asian, European, Native American, Oceanian, South Asian, and West Asian. We then used RFMix v2.03^33^ to assign each of the 20,727 windows across the phased genomes to one of these eight ancestry clusters (for all individuals in the UK Biobank).

These local ancestry assignments were subsequently used with PLINK2 as local covariates in the GWAS for the admixed individuals for SNPs within those respective windows. PLINK2 allows for the direct input of the RFMix output (the MSP file, which contains the most likely subpopulation assignment per conditional random field [CRF] point) as local covariates using the “local-cov”, “local-psam”, and “local-haps” flags, the “local-cats0=n” flag (where n is the number of assignments), and the “local-pos-cols=2,1,2,7” flag (for a typical RFMix MSP output file - see https://www.cog-genomics.org/plink/2.0/assoc).

#### Variant Quality Control and Metadata Generation

For quality control (QC), In total, we ensured that variant-level missingness was less than 10%, that the *p*-value for the Hardy-Weinberg equilibrium test (computed within unrelated individuals of white British ancestry) was greater than 10^−15^, and that the variant was uniquely represented (the “CHROM:POS:REF:ALT” variant string was uniquely identified) in the PLINK dataset file. In total, we removed 195,920 variants that failed to meet all of these criteria, except for 134 variants on the Y chromosome.

For the remainder, we used Variant Effect Predictor (VEP)^34^ to annotate the most severe consequence, the gene symbol, and HGVSp of each variant in the UK Biobank exome and array data. We calculated minor allele frequencies using PLINK. MPC^11^ values (variant-level) and pLI gene memberships^12^ were annotated from source. To determine LD independence criteria, we used PLINK’s --indep-pairwise function with a window size of 1000kb, a step size of 1, and an r^2^ threshold of 0.1 on those variants that pass QC. As our analyses focused on PTVs and PAVs, we then performed this same LD independence analysis on only these, overriding assignments in the first analysis if necessary. We provide these essential metadata, which are necessary for MRP, in exome (https://biobankengine.stanford.edu/static/ukb_exm_oqfe-consequence_wb_maf_gene_ld_indep_mpc_pli.tsv.gz) and array (https://biobankengine.stanford.edu/static/ukb_cal-consequence_wb_maf_gene_ld_indep_mpc_pli.tsv.gz) tables, available for direct download via the Global Biobank Engine.^35^

#### Applications

For exome applications, we chose variants with MAF ≤ 1% and that were LD-independent according to the criteria mentioned above. For quantitative traits, we removed variants whose regression effect size had standard error greater than 100, and for binary traits, we removed variants whose regression effect size had standard error greater than 0.2. For array applications, we chose variants with MAF ≤ 1% and removed variants whose regression effect size had standard error greater than 0.2. While MRP is capable of handling all variant types (e.g. proximal coding and intronic variants), we included only protein-altering variants (PAVs) and protein-truncating variants (PTVs) in both exome and array analyses (exome data features many more PAVs and thus potential for power gain; **Table S1**; **Figure S2**). These sets respectively contain the following consequence annotations:

- PAVs: protein_altering_variant, inframe_deletion, inframe_insertion, splice_region_variant, start_retained_variant, stop_retained_variant, missense_variant
- PTVs: frameshift_variant, splice_acceptor_variant, splice_donor_variant, stop_gained, start_lost, stop_lost

For both quantitative and binary traits, PTVs were assigned a σ (standard deviation of prior on effect size) of 0.2, whereas PAVs were assigned a σ value of 0.05. We also incorporated MPC and pLI deleteriousness metrics into our exome analyses. For those PTVs with a pLI of > 0.8, we increased σ to 0.5, and for those PAVs with an MPC ≥ 1, we set σ = 0.05 x MPC. These adjustments serve to further granularize and weight MRP results in biologically meaningful ways (**Table S3**; **Figure S4**). For the exome meta-analysis, we assumed a similar effects model across studies and an independent effects model across variants.

We also studied how the application of MRP to multiple phenotypes together would potentially boost power to detect rare-variant associations. We calculated pairwise genetic correlation between 35 biomarker phenotypes^17^ using LD score regression,^25^ and then used the hclust algorithm in the R stats package^36^ to generate phenotype clusters. For each of these clusters, using the array data, we performed MRP in the multi-phenotype setting.

## Results

### Simulations

To study the behavior of MRP going from a single phenotype to multiple phenotypes, we conducted a simulation study where we assumed an allelic architecture consistent to that discovered for *APOC3* in relation to triglycerides, low-density lipoprotein cholesterol (LDL-C), and high-density lipoprotein cholesterol (HDL-C).^37–39^ We simulated three continuous phenotypes with a total correlation consistent with that observed for triglycerides, LDL-C, and HDL-C. Furthermore, we introduced effects to four variants consistent with the effects observed in four PTVs (approximately 0.35 standard deviations away from the population mean) and to another four variants consistent with the effects observed for missense variants (approximately 0.2 standard deviations away from the population mean) all with minor allele frequency of 0.05%. The PTV group of variants had the same effects, whereas out of the missense variants, half had positive and the other half had negative effect sizes. The correlation of effects between the group of phenotypes was set to be directionally consistent with the direction of genetic effects observed for lipid phenotypes and PTVs in *APOC3*, i.e. proportional effects for triglycerides and LDL-C, and inversely proportional for LDL-C and HDL-C, and triglycerides and HDL-C. We simulated 1000 genes where 50 of the genes contained non-zero effects on the multivariate phenotype. Given we know which of the 1000 genes contained non-zero effects, we could compute the true positive rates and false positive rates for a given BF threshold. We analyze the data as follows: i) single-variant and single-phenotype, ii) multiple variants and single-phenotype, iii) single-variant and multiple-phenotypes, and iv) multiple-variants and multiple-phenotypes (**Figure 2A**). We find that in some scenarios, analyzing multiple-variants and multiple-phenotypes jointly improved the ability to detect signals.

**Figure 2A.**
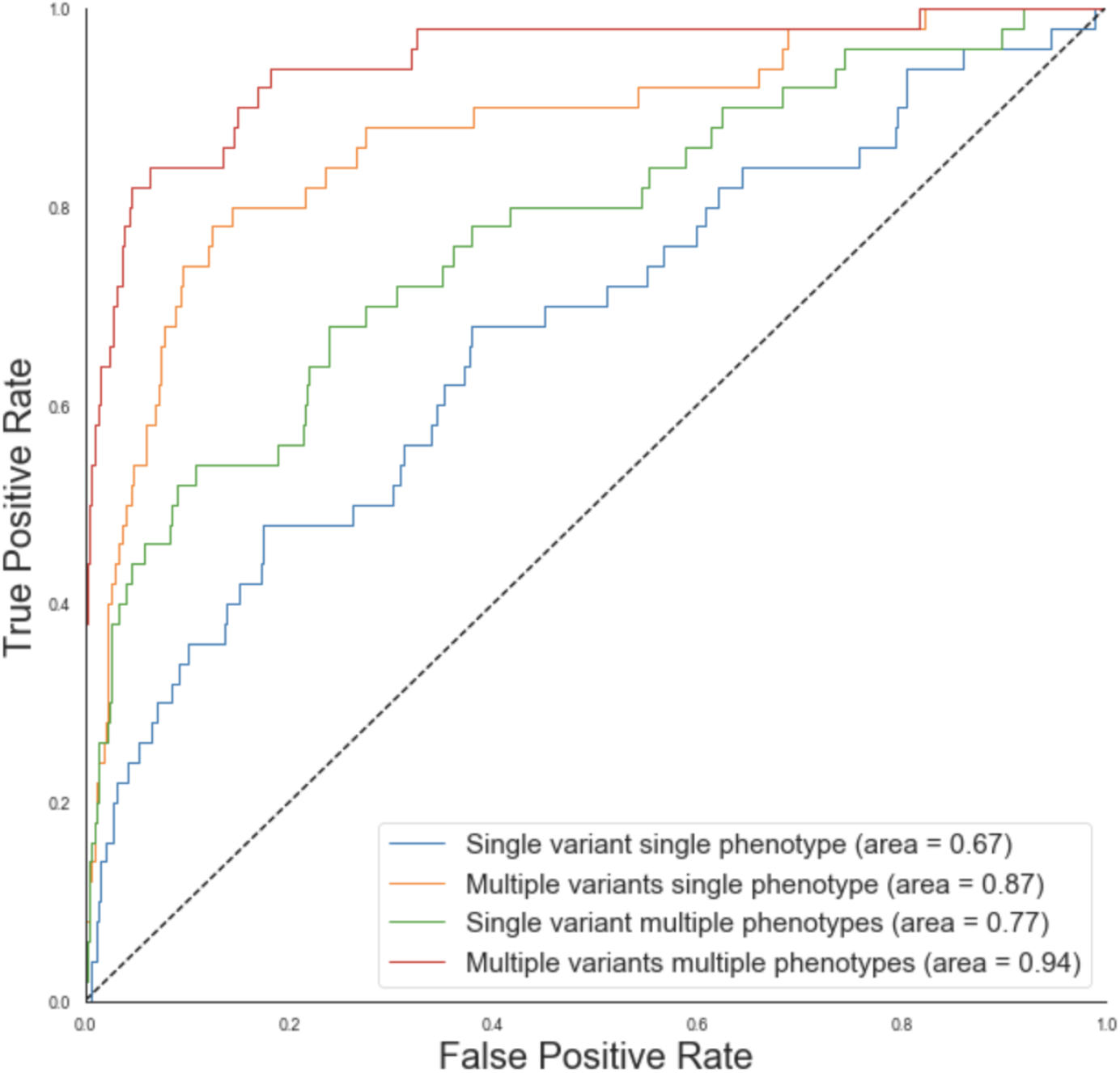
From single-variant and single-phenotype to multiple-variant and multiple-phenotype gene discovery. ROC curves for detecting simulated gene association to any of the phenotypes using single variant/single phenotype association (blue) to multiple-variant and multiple-phenotype association (red).

### Exome single-phenotype meta-analyses

MRP was used to perform exome meta-analysis on 2019 traits across six UK Biobank populations as described in **Methods**. Among the best powered and represented traits were a set of 35 biomarkers, the focus of a previous publication^17^. We see the number of log_10_ BF ≥ 5 genes increasing from a single-population to a meta-analysis setting. Since we expect that the meta-analysis over different ancestries cannot be more confounded than the analysis of a single ancestry, we interpret the increase in the number of genes as an increase in the statistical power to detect rare-variant associations (**Figure 2B**).

**Figure 2B.**
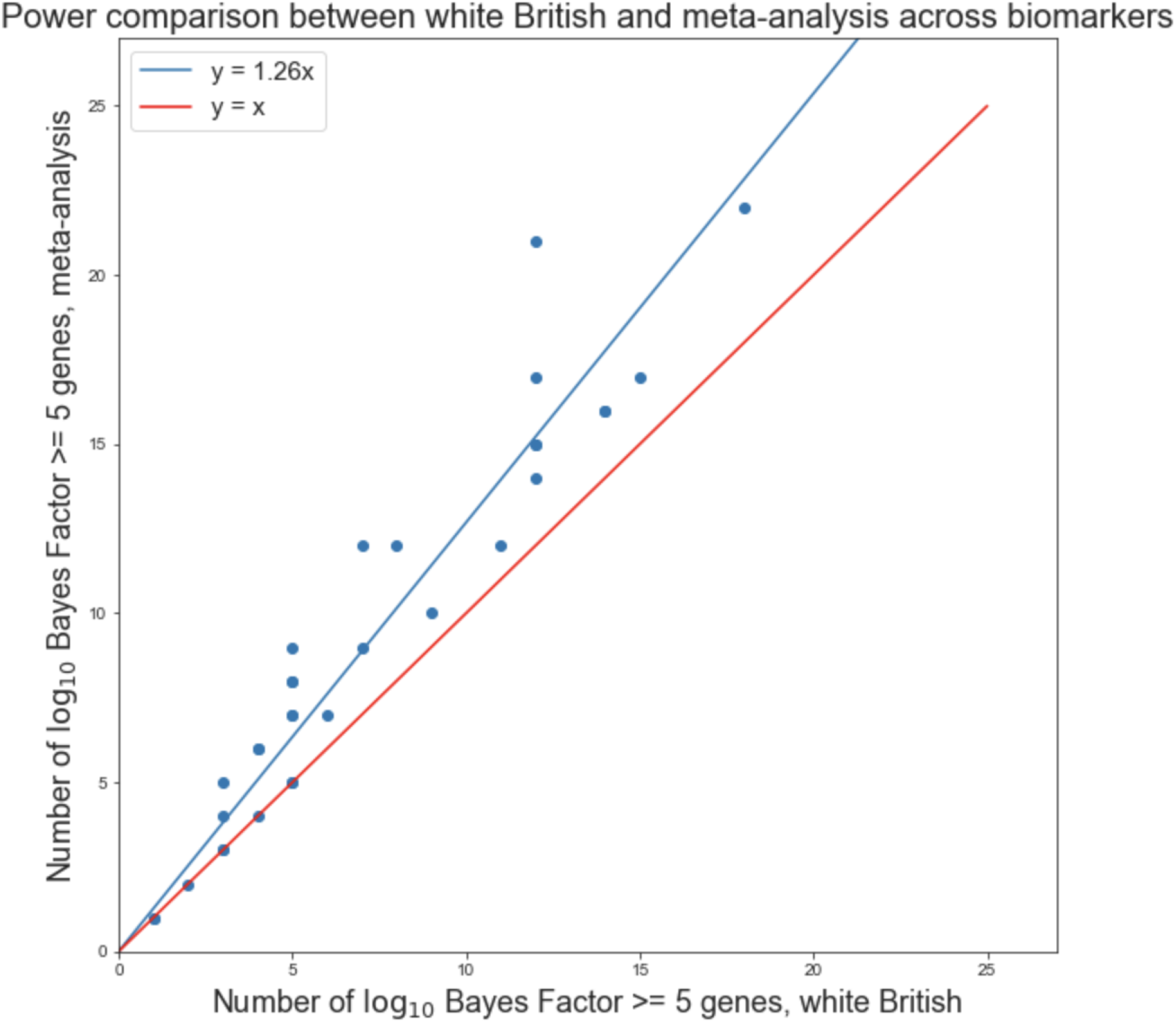
From single to multiple populations. Scatterplot showing number of genes with log_10_ BF ≥ 5 for white British population only (x-axis) versus meta-analysis (y-axis) across 35 biomarkers. Assuming that BFs are correctly calibrated in both analyses and that meta-analysis is not inflated compared to white British-only MRP, suggests a ∼26% increase in power when incorporating summary statistics across multiple populations.

We categorize these biomarkers into six categories as in Sinnott-Armstrong et. al.^17^ (Bone and Joint, Cardiovascular, Diabetes, Hormone, Liver, and Renal - **Figure 3**) and we recover several known gene-trait associations and discover several others.

**Figure 3.**
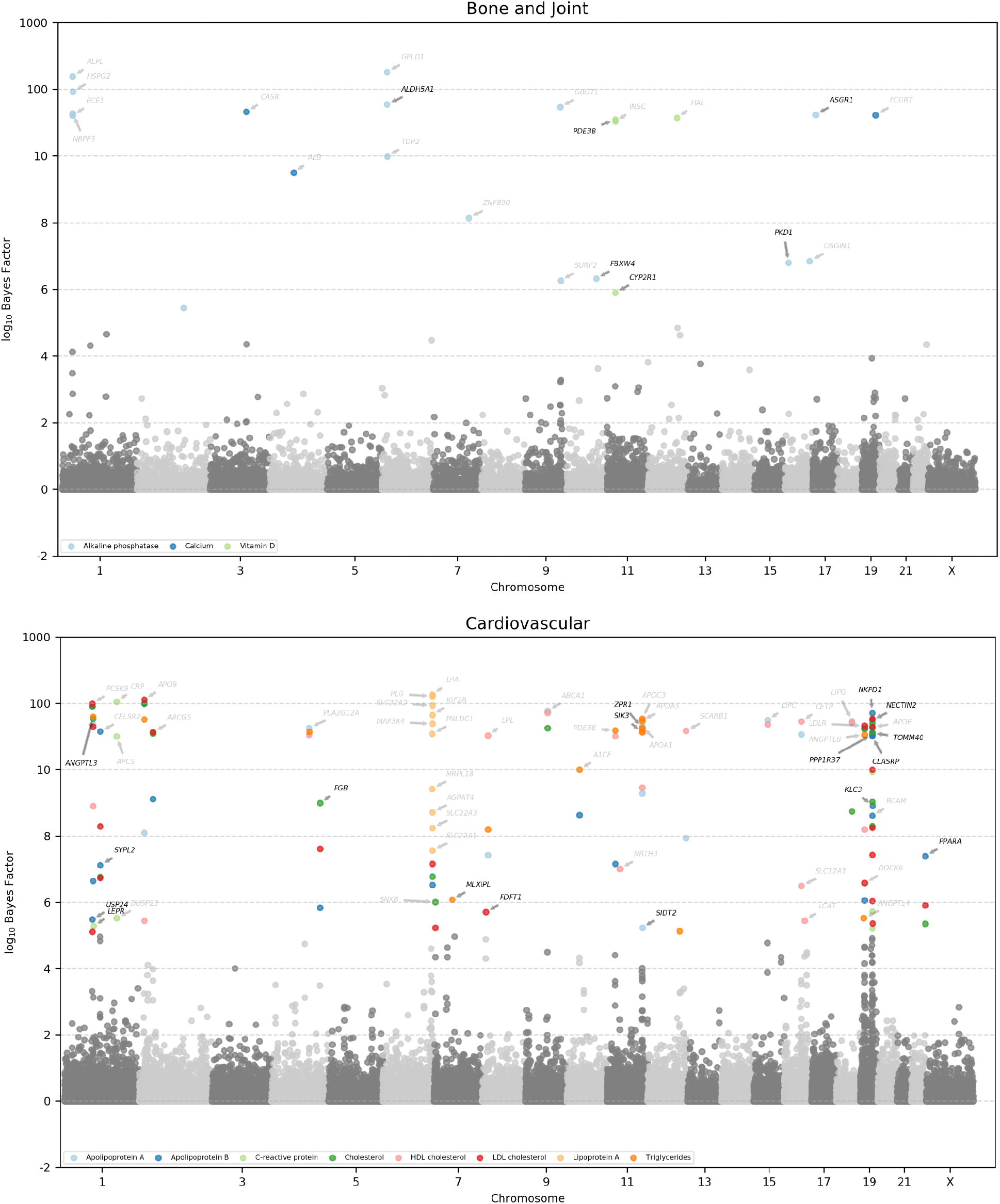
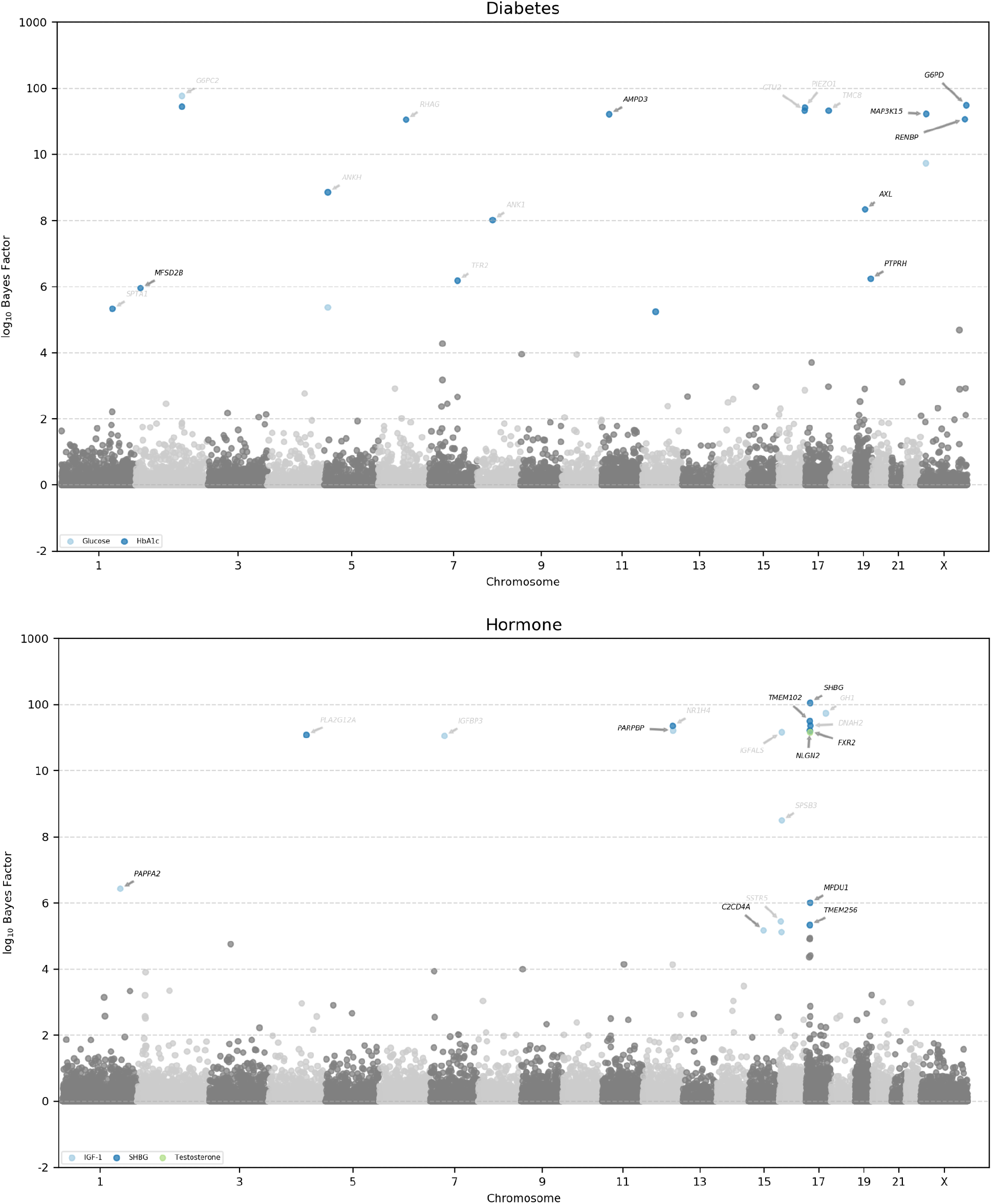
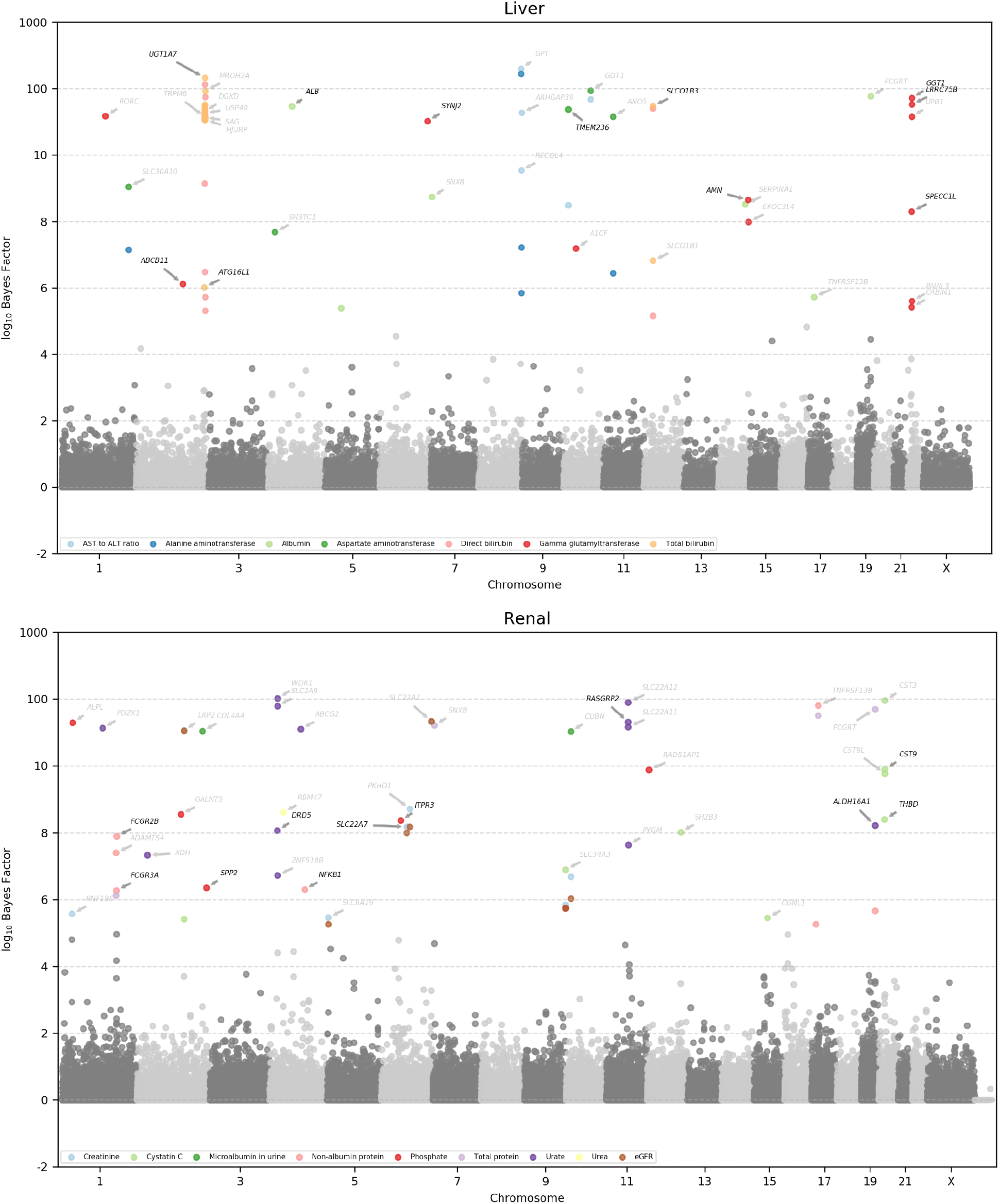
Manhattan plots showing log_10_ BF under an independent effects variant model amongst protein-altering variants for 6 categories across 35 biomarkers. Scale is logarithmic after log_10_ BF ≥ 10. Genes found in Sinnott-Armstrong, et.al.^17^ are annotated in grey, whereas the other genes are annotated in black.

Among the “Bone and Joint” biomarkers (alkaline phosphatase, calcium, and vitamin D), we recover associations between *CASR* and calcium^40^ and *HAL* and vitamin D^41^. As compared to results from array data as found in Sinnott-Armstrong, et. al.^17^, we also recover exome-specific associations between *ALDH5A1* and alkaline phosphatase^42^ and *PDE3B* and vitamin D^41^.

For the “Cardiovascular” phenotypes (apolipoprotein A, apolipoprotein B, C-reactive protein, total cholesterol, HDL cholesterol, LDL cholesterol, lipoprotein A, and triglycerides), MRP recovers array associations between: *PLG, LPA*, and lipoprotein A^43^; *APOC3* and triglycerides^44^; *ANGPTL3* and triglycerides^45^; *APOB* and apolipoprotein B^46^ and LDL cholesterol^47^; *ABCA1* and apolipoprotein A^44^ and HDL cholesterol^47^; *PCSK9* and total cholesterol^48^; and *CRP* and C-reactive protein^49^. Exome-only signals recover associations such as between *ZPR1*^*50*^ and *SIK3*^*44*^ and triglycerides.

In the two diabetes-related phenotypes (glucose and HbA1c), we recover associations between *G6PC2* and glucose^48^ as well as *PIEZO1* and HbA1c^51^ and an additional exome association between *G6PD* and HbA1c^51^. Hormonal recoveries include those between *SHBG* and SHBG and testosterone levels and *GH1* and IGF-1 levels.

MRP applied to liver-related phenotypes recover known associations between: *UGT* genes and bilirubins^52^; *GOT1* and aspartate aminotransferase^53^; *FCGRT* and albumin^42^; and *GPT* and AST-ALT ratio^49^. in the exome sequencing, we additionally recover associations between *GGT1* and gamma glutamyltransferase^54^, *TMEM236* and aspartate aminotransferase^42^, and *SLCO1B3* and bilirubin.^55^

The renal traits similarly feature a mix of array recoveries and exome discoveries. We recover signal between: *SLC22A2* and creatinine^56^; *CST3* and Cystatin C^46^; *COL4A4* and microalbumin^57^; *TNFRSF13B* and non-albumin protein^42^; *FCGRT* and total protein; *WDR1, RASGRP2, DRD5*, and urate^58,59^; and *LRP2* and eGFR levels^60^. We additionally discover novel gene-trait associations (not found in the NHGRI-EBI catalog or Open Targets Genetics) across these biomarker categories, including: *GLPD1* and alkaline phosphatase; *NKPD1* and apolipoprotein B; *RENBP/MAP3K15* and Hba1c; *PARPBP* and IGF-1; *NLGN2* and SHBG; *ALB* and albumin; *ALPL* and phosphate; *RBM47* and urea; *ALDH16A1* and urate; *THBD* and Cystatin C; *ITPR3* and phosphate; *SLC22A7* and creatinine; and *FCGR2B* and non-albumin protein.

For the 2,019 traits for which MRP was performed, there were also a considerable number of associations found amongst non-biomarker traits. We found associations between: *TUBB1* and platelet distribution width and mean platelet volume^61^; *IL17RA* and monocyte count and percentage^61^; *OCA2*/*MC1R* and skin color/hair color^62–65^; *IQGAP2* and mean platelet volume^61^; *SLC24A5, HERC2, TCF25, TYR* and skin color^66^; *SH2B3, JAK2* and platelet crit^67^ and count^68^; *KALRN* and mean platelet volume^61^; *HBB* and mean corpuscular volume^69^, mean corpuscular hemoglobin^70^, and red blood cell count^71^; and *CXCR2* and neutrophil count^61^.

We have published the full set of associations (log_10_ BF ≥ 5) from an independent effects model amongst PAVs, from a similar effects model amongst PAVs, as well as from a similar effects model amongst PTVs on the Global Biobank Engine for exomes (https://biobankengine.stanford.edu/RIVAS_HG38/mrpgene/all) and array data (https://biobankengine.stanford.edu/RIVAS_HG19/mrpgene/all).

MRP was implemented using Python (dependencies: pandas v1.1.5, numpy v1.16.4, rpy2 v3.0.4, scipy v1.3.0). The requirements, code, and metadata files can be found at https://github.com/rivas-lab/mrp.

### Array single-population multi-phenotype analyses

In order to demonstrate the effectiveness of MRP to boost signal in a multi-phenotype context, we used LD-score regression^25^ to determine genetic correlations between the 35 biomarker traits (**Figure S5**) that were a focus of a previous paper^17^. This correlation matrix was then used for hierarchical clustering followed by dynamic tree cutting, which formed four clusters of between seven and ten traits each (**Figure 4A**). We generated the correlation plots as shown in **Figure 4B**.

**Figure 4A.**
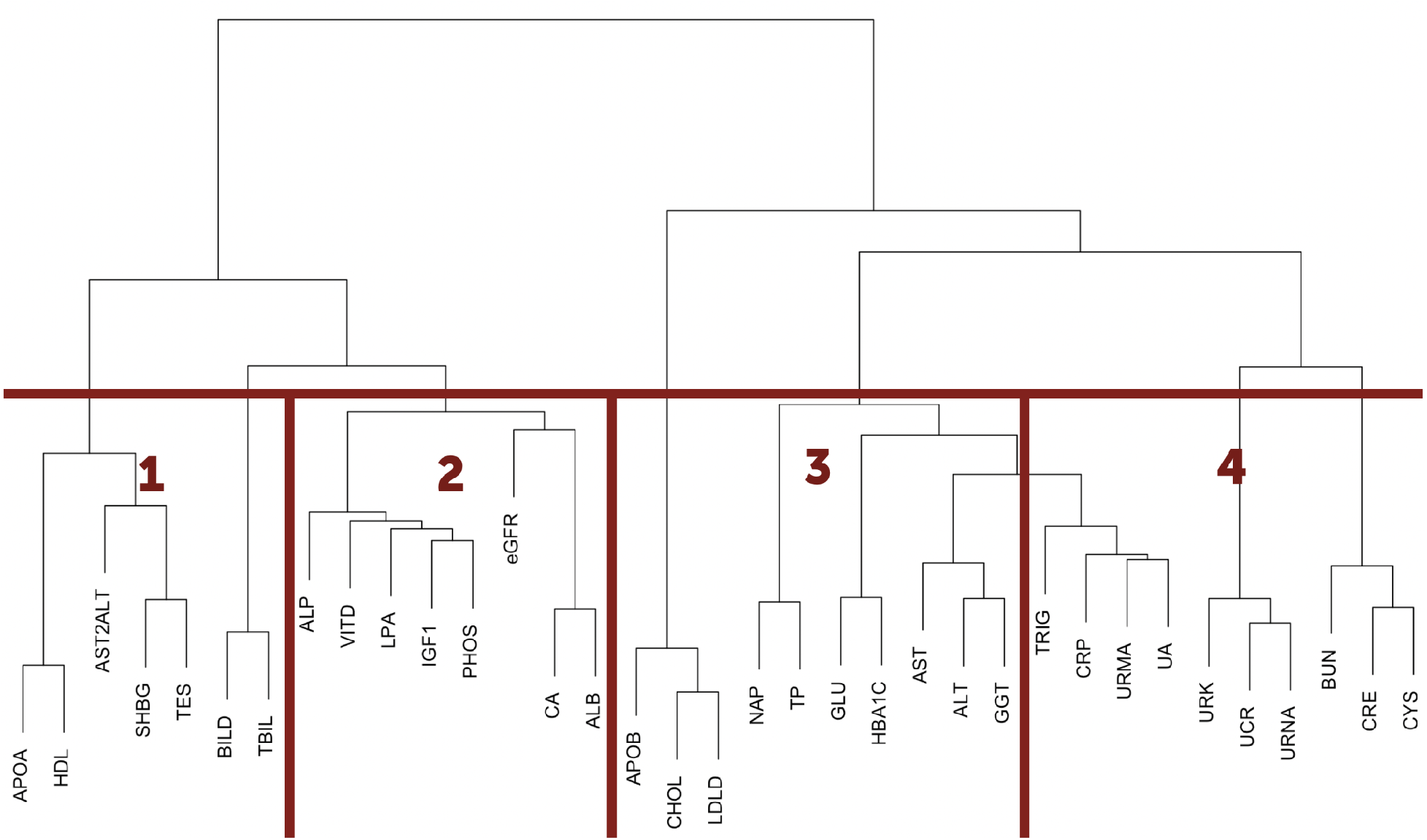
Hierarchical clustering dendrogram. Based on genetic correlationderived from an LD-score regression-based distance matrix between 35 biomarker traits.

**Figure 4B.**
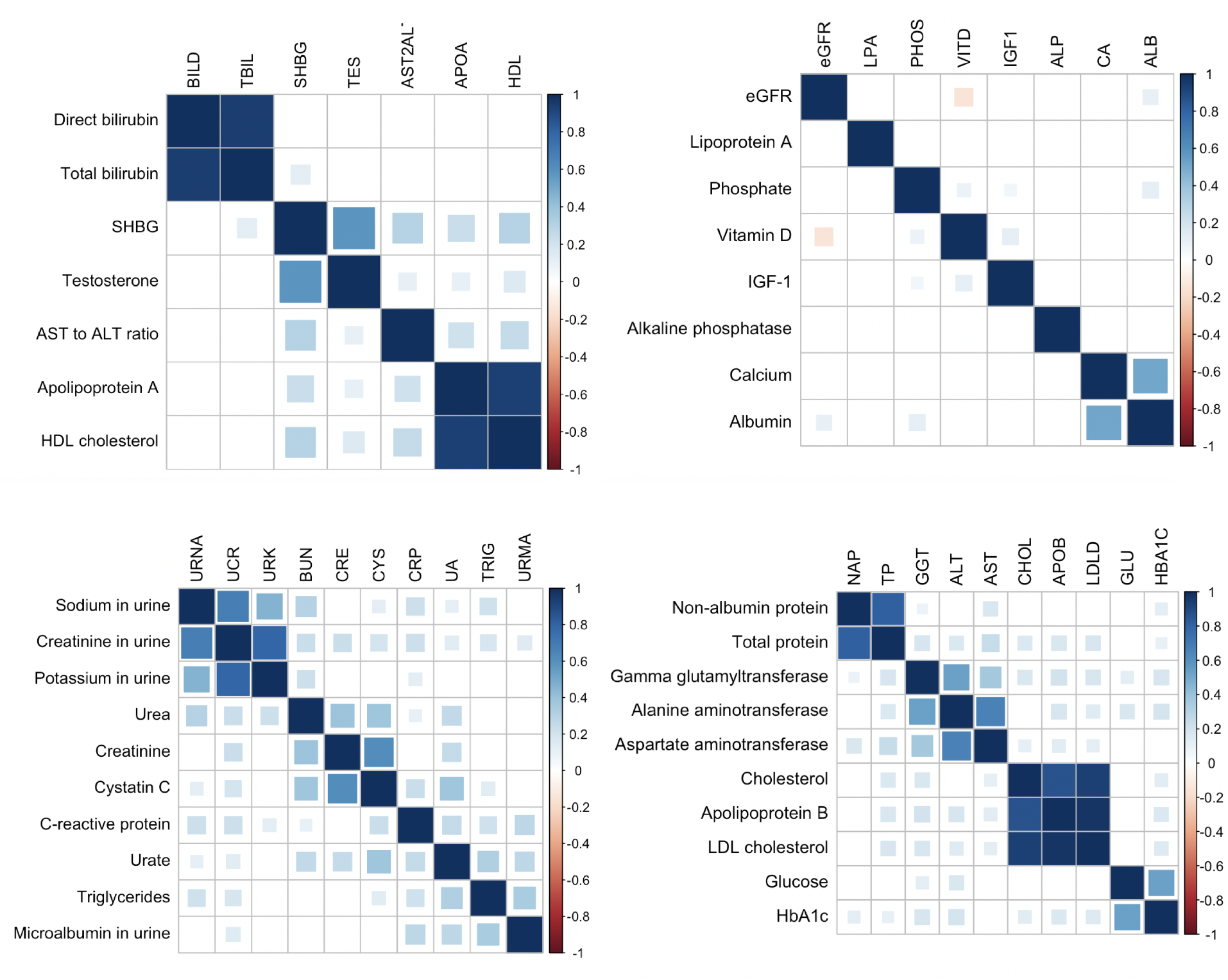
LD-score regression-based genetic correlation plots of candidate clusters. Derived from the dendrogram in **Figure 4A** using a dynamic tree cutting algorithm.

Multi-phenotype MRP results in several substantial power gains throughout the four clusters; one of these clusters is highlighted in **Figure 4C**. As compared to the maximum log_10_ BF from the constituent phenotypes, the multi-phenotype analysis generally fares comparably, while also highlighting clear targets. We found evidence for association between rare coding variants in several genes and the clusters above; *TNFRSF13B* (log_10_ BF_multi-trait_ = 204.5, max[log_10_ BF_single-trait_] = 141.0), *APOB* (log_10_ BF_multi-trait_ = 197.9, max[log_10_ BF_single-trait_] = 128.0), *SNX8* (log_10_ BF_multi-trait_ = 96.0, max[log_10_ BF_single-trait_] = 43.8) receive a boost in log_10_ BF of over 50 units for cluster 1 (Alanine aminotransferase, Aspartate aminotransferase, Gamma glutamyltransferase, Glucose, HbA1c, Total protein, Apolipoprotein B, Cholesterol, LDL cholesterol, and Non-albumin protein). Several other genes that are clearly below 5 (in log_10_ BF) in the single-trait settings become above 5 in the joint setting (e.g., *G6PC*; log_10_ BF_multi-trait_ = 5.3, max[log_10_ BF_single-trait_] = 1.3). The *G6PC* gene provides instructions for making the glucose 6-phosphatase enzyme, found on the membrane of the endoplasmic reticulum. The enzyme is expressed in active form in the liver, kidneys, and intestines, and is the main regulator of glucose production in the liver; given the traits included in cluster 1, the increase in power may be biologically relevant^72^. These results demonstrate that MRP can identify biologically meaningful targets that may be missed by standard GWAS approaches.

**Figure 4C.**
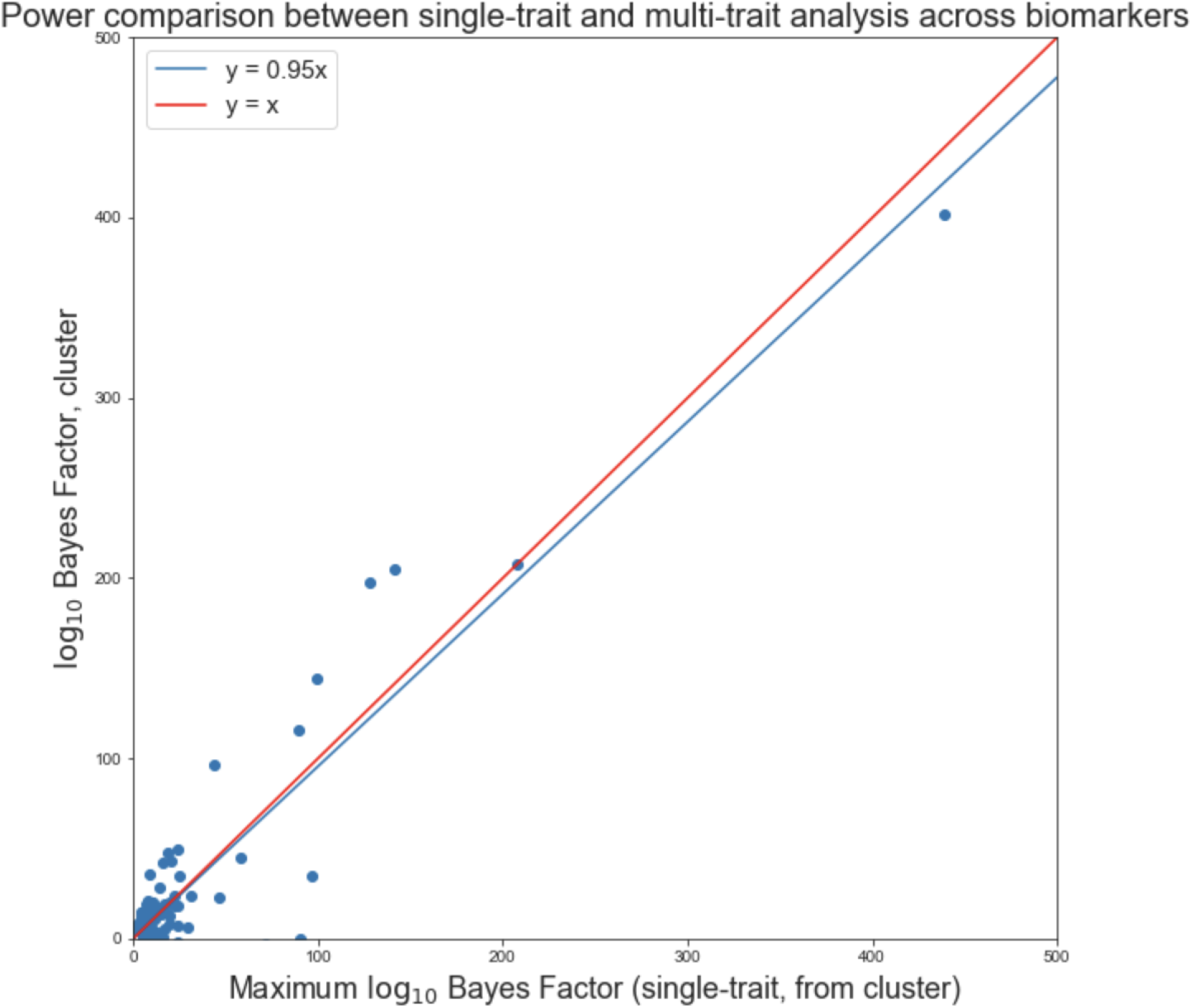
Cluster vs. single-trait power analysis. Power comparison of genes with log_10_ BF ≥ 5 in either i) any of the single-trait analyses of the traits within the cluster or ii) the multi-trait analysis, for a cluster of biomarkers (Alanine aminotransferase, Aspartate aminotransferase, Gamma glutamyltransferase, Glucose, HbA1c, Total protein, Apolipoprotein B, Cholesterol, LDL cholesterol, and Non-albumin protein). *x*-axis depicts the maximum log_10_ BF of the gene amongst any of the constituent single-trait analyses, and *y*-axis depicts the multi-trait result. Multi-trait analyses roughly equal the highest-powered single-trait analyses, while also substantially boosting signal in some genes.

## Discussion

In this study, we developed MRP, a Bayesian model comparison approach that shares information across variants, phenotypes, and studies to identify rare variant associations. We used simulations to verify that jointly considering both variants and phenotypes can improve the ability to detect associations. We also applied the MRP model comparison framework in a meta-analysis setting to exome summary statistics across the UK Biobank, identifying strong evidence for the previously described associations between, for example, *HAL* and vitamin D^41^, and discovering several novel associations, such as between *GLPD1* and alkaline phosphatase. We made the full results set available on the Global Biobank Engine (https://biobankengine.stanford.edu/)^35^. We also leveraged MRP to boost signal in a multi-phenotype setting using the array data (which has many more samples than the exome data), finding genes such as *G6PC* that do not come up in the single-trait context but show strong evidence in the joint analysis. These results demonstrate the ability of the MRP model comparison approach to leverage information across multiple phenotypes and variants to discover rare variant associations.

As genetic data linked to high-dimensional phenotype data is increasingly being made available through biobanks, health systems, and research programs, there is a large need for statistical approaches that can leverage information across different genetic variants, phenotypes, and studies to make strong inferences about disease-associated genes. The approach presented here relies only on summary statistics from marginal association analyses, which can be shared with less privacy concerns compared to raw genotype and phenotype data. Combining joint analysis of variants and phenotypes with meta-analysis across studies offers new opportunities to identify gene-disease associations.

## Author Contributions

M.A.R. and M.P. designed the method and derived all analytical calculations. G.R.V. implemented the method and analyzed the data. G.R.V., M.A.R., M.P., and C.D. wrote the manuscript. G.R.V., M.A.R., M.P., C.C.A.S., C.D., Y.T., M.A., T.P., and H.M. provided quality control analysis, figure edits, and revisions to the manuscript. A.G.I. also provided revisions and feedback on local ancestry-corrected GWAS. C.D.B. and M.J.D. provided critical feedback on methodology.

## Acknowledgements and Funding

This research was conducted using the UK Biobank Resource under application number 24983, “Generating effective therapeutic hypotheses from genomic and hospital linkage data” (http://www.ukbiobank.ac.uk/wp-content/uploads/2017/06/24983-Dr-Manuel-Rivas.pdf). Based on the information provided in protocol 44532, the Stanford IRB has determined that the research does not involve human subjects as defined in 45 CFR 46.102(f) or 21 CFR 50.3(g). All participants in the UK Biobank study provided written informed consent (more information is available at https://www.ukbiobank.ac.uk/2018/02/gdpr/). Statin adjustment analyses were further conducted via UK Biobank application 7089 using a protocol approved by the Partners HealthCare Institutional Review Board. We thank all the participants in the UK Biobank. We thank members of the Rivas lab for their feedback. M.A.R. is in part supported by the NHGRI of the NIH under award R01HG010140 (M.A.R.) and an NIH Center for Multi- and Trans-ethnic Mapping of Mendelian and Complex Diseases grant (5U01 HG009080). G.R.V. is supported by the National Library of Medicine (NLM) T15 Continuing Education Training Grant. The content is solely the responsibility of the authors and does not necessarily represent the official views of the NIH. Some of the computing for this project was performed on the Sherlock cluster at Stanford University. We would like to thank Stanford University and the Stanford Research Computing Center for providing computational resources and support that contributed to these research results.

## Supplementary Materials

**Table S1.** Genes with considerable power gain in exome data as compared to array data.

| Trait | gene | Number of PAVs, array | $\log_{10}BF$, array | Number of PAVs, exome | $\log_{10}BF$, exome | $\log_{10}BF$ Difference |
| --- | --- | --- | --- | --- | --- | --- |
| Total bilirubin | *UGT1A7* | 5 | 1.2 | 247 | 213 | 211.8 |
| Direct bilirubin | *UGT1A7* | 5 | 0.6 | 228 | 133 | 132.4 |
| Lipoprotein A | *PLG* | 57 | 38.9 | 583 | 165 | 126.1 |
| SHBG | *SHBG* | 7 | 2.7 | 284 | 114 | 111.3 |
| LDL cholesterol | *PCSK9* | 94 | 4.0 | 759 | 99 | 95.0 |
| Total bilirubin | *MROH2A* | 33 | 4.6 | 1649 | 85.8 | 81.2 |
| Apolipoprotein B | *PCSK9* | 94 | 3.1 | 756 | 80.7 | 77.6 |
| Cholesterol | *PCSK9* | 94 | 4.0 | 760 | 80.9 | 76.9 |
| IGF-1 | *GH1* | 5 | 2.1 | 301 | 55.1 | 53.0 |
| Direct bilirubin | *MROH2A* | 33 | 2.9 | 1497 | 55.7 | 52.8 |
| Gamma glutamyltransferase | *GGT1* | 5 | 0.008 | 545 | 52.1 | 52.1 |
| Triglycerides | *ANGPTL3* | 7 | -0.02 | 337 | 39.9 | 39.9 |
| Cholesterol | *ANGPTL3* | 7 | -0.6 | 337 | 34.3 | 34.9 |
| Cholesterol | *APC* | 1409 | -34.7 | 1882 | -0.5 | 34.2 |
| LDL cholesterol | *APC* | 1410 | -33.7 | 1882 | -0.5 | 33.2 |
| Apolipoprotein B | *APC* | 1405 | -32.7 | 1876 | -0.7 | 32.0 |
| Total bilirubin | *UGT1A5* | 12 | 2.0 | 225 | 33 | 31.0 |
| Albumin | *APC* | 1366 | -31.6 | 1807 | -1.2 | 30.4 |
| Vitamin D | *APC* | 1379 | -29.8 | 1828 | 0.2 | 30.0 |
| Creatinine | *APC* | 1411 | -31.4 | 1883 | -1.9 | 29.5 |

**Figure S2.**
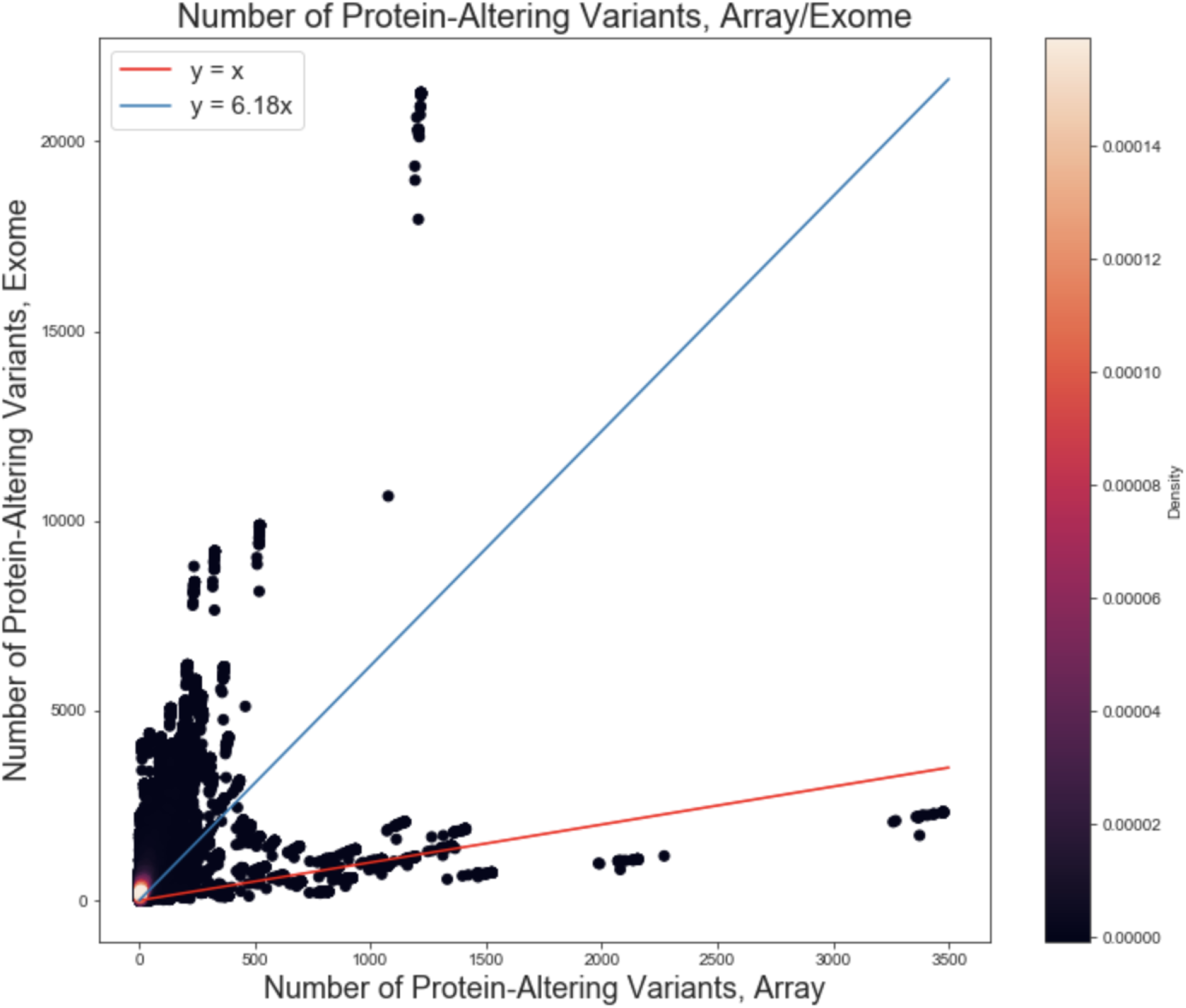
From array to exome. Scatterplot showing the increase in number of protein-altering variants in genes used in the analysis when comparing array (*x*-axis) to exome (*y*-axis) data. Data is taken from MRP calculations across 35 biomarker traits within the UK Biobank.

**Table S3.** Power comparison between variant annotation-based MRP and MPC/pLI-augmented MRP analyses across 35 biomarkers. We see considerable gains in power in several gene/trait combinations.

| Trait | Gene | Number of PAVs | $\log_{10}$BF without MPC | Number of MPC-augmented PAVs | Number of pLI-augmented PAVs | $\log_{10}$BF with MPC | $\log_{10}$BF Difference |
| --- | --- | --- | --- | --- | --- | --- | --- |
| Alkaline phosphatase | *ALPL* | 198 | 126 | 93 | 0 | 160 | 34 |
| Lipoprotein A | *LPA* | 512 | 109 | 20 | 0 | 114 | 5 |
| Apolipoprotein A | *APOA1* | 102 | 11.7 | 30 | 0 | 15.7 | 4 |
| HDL cholesterol | *APOA1* | 103 | 9.36 | 30 | 0 | 13.2 | 3.84 |
| Aspartate aminotransferase | *SLC30A10* | 112 | 3.76 | 50 | 6 | 7.2 | 3.44 |
| Phosphate | *ALPL* | 192 | 10.9 | 91 | 0 | 14.3 | 3.4 |
| Lipoprotein A | *IGF2R* | 763 | 29.8 | 153 | 27 | 33.1 | 3.3 |
| HDL cholesterol | *SCARB1* | 220 | 5.45 | 66 | 0 | 8.29 | 2.84 |
| Apolipoprotein B | *APOE* | 142 | 5.48 | 60 | 0 | 8.27 | 2.79 |
| Alanine aminotransferase | *SLC30A10* | 112 | 2.94 | 50 | 6 | 5.56 | 2.62 |

**Figure S4.**
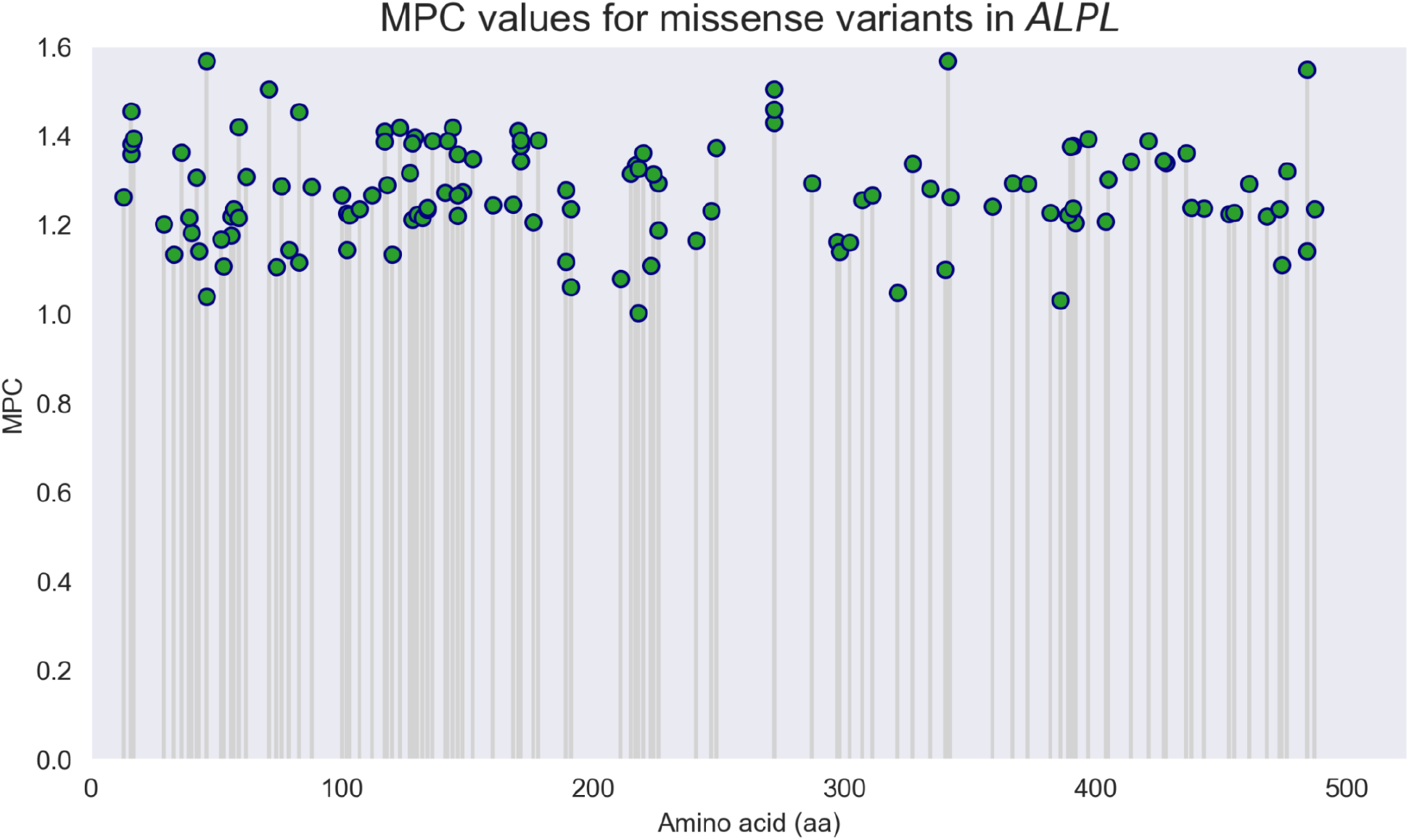
*ALPL* gene plot. Gene plot showing variants for which MPC pathogenicity information was incorporated, resulting in a power gain for *ALPL* gene that encodes alkaline phosphatase; for the Alkaline phosphatase phenotype, the incorporation of this information resulted in a log_10_BF gain of 34 (**Table S3**).

**Figure S5.**
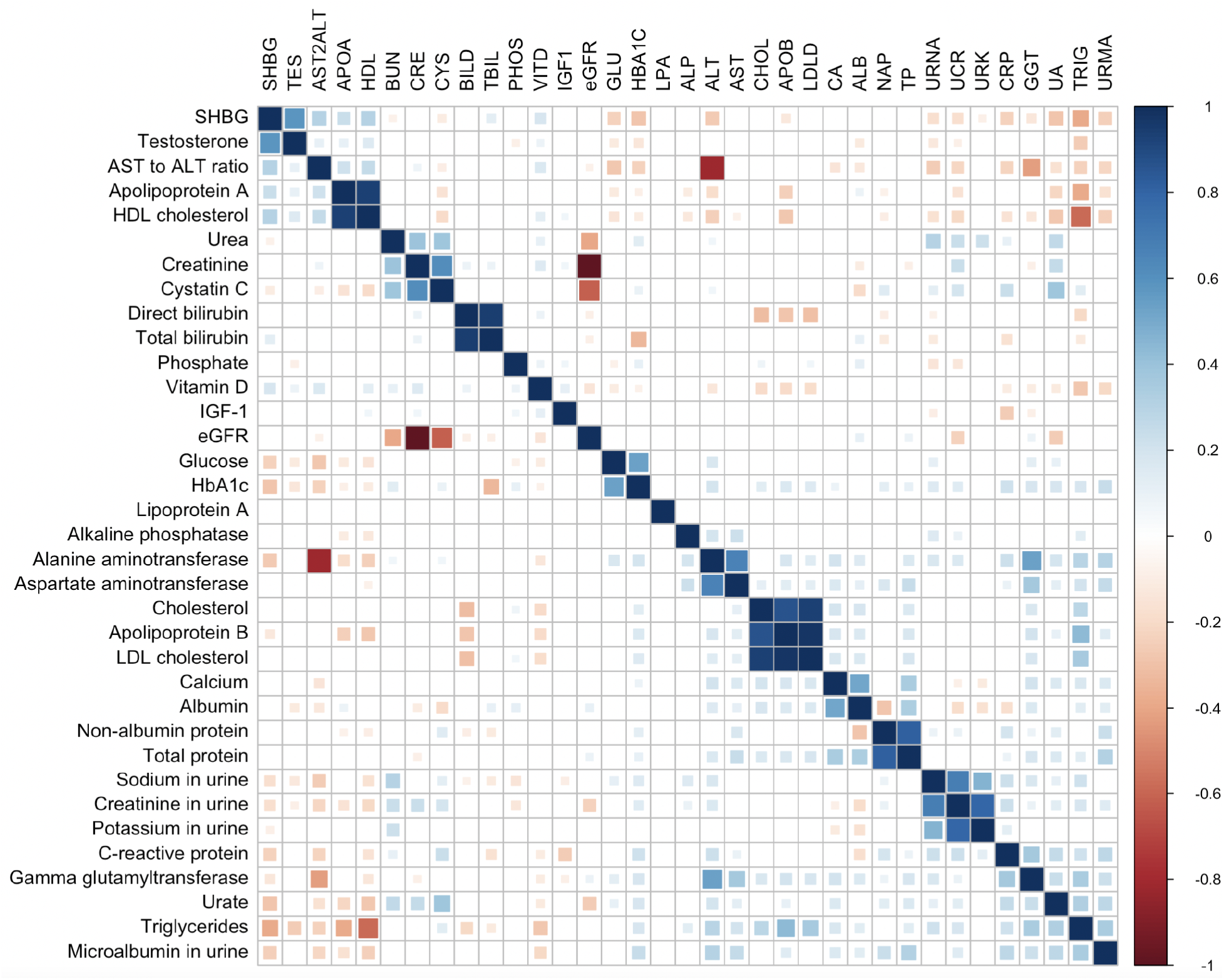
LD-score regression-based genetic correlation plots of all 35 biomarkers included in the multi-trait analyses. The traits are ordered by hierarchical clustering.

## Notes

https://biobankengine.stanford.edu/RIVAS_HG38/mrpgene/all

### Competing Interest Statement

M.A.R. is on the SAB of 54Gene, Related Sciences and scientific founder of Broadwing Bio and has advised BioMarin, Third Rock Ventures and MazeTx. C.D.B. is the Owner and President of C.D.B. Consulting, LTD. and also a Director at EdenRoc Sciences, LLC and Etalon DX, founder of Arc Bio LLC (formerly IdentifyGenomics LLC and BigData Bio LLC), and an SAB member of Imprimed, FaunaBio, Columbia Care, and Digitalis Ventures. He is also a Venture Partner at F-Prime Capital Partners. M.J.D. is a founder of MazeTx.

### Summary of Updates

This revision has been updated to include results from new exome data, a productionization of the python package, and a visualization tool in the Global Biobank Engine. We include an Acknowledgements and Author Contributions statement as well.

