## Supplemental Note for "Bayesian model comparison for rare variant association studies"

July 7, 2021

#### 1 MRP model comparison for association testing

We consider the multivariate linear regression model

$$\underset{(N\times K)}{\mathrm{Y}} = \underset{(N\times K)}{\mathbf{\Psi}} + \underset{(N\times M)}{\mathrm{X}}\underset{(M\times K)}{\mathrm{B}} + \underset{(N\times K)}{\mathrm{E}},$$

where the matrices $\mathrm{Y} = [y_{ik}]$, $\mathrm{X} = [x_{im}]$, $\mathrm{B} = [\beta_{mk}]$ and $\mathrm{E} = [e_{ik}]$ describe the phenotype values ($y_{ik}$), copies of minor allele ($x_{im}$), variant-phenotype effects ($\beta_{mk}$), and residual errors ($e_{ik}$), for individual $i$, phenotype $k$, and variant $m$. We assume that each phenotype has been transformed to a standard normal distribution and that the columns of X have been centered, which means that the estimate for the intercept term $\mathbf{\Psi}$ is 0 and independent of the estimate of B. We use vectorized notation where the rows of B form vector $\boldsymbol{\beta} = (\boldsymbol{\beta}_1, \ldots, \boldsymbol{\beta}_M)^\intercal$ of length $MK$.

We define the MRP model comparison as a Bayes factor (BF) between the alternative model, where at least one variant affects at least one phenotype, and the null model, where all variant-phenotype effects are zero. BF is the ratio of the marginal likelihoods for these two models:

$$\mathrm{BF} = \frac{\int_{\boldsymbol{\beta}} p\left(\mathrm{Data}|\boldsymbol{\beta}\right) p\left(\boldsymbol{\beta}|\mathrm{ALT}\right) d\boldsymbol{\beta}}{\int_{\boldsymbol{\beta}} p\left(\mathrm{Data}|\boldsymbol{\beta}\right) p\left(\boldsymbol{\beta}|\mathrm{NULL}\right) d\boldsymbol{\beta}},$$

where Data can correspond either to the effect size estimates $\widehat{\boldsymbol{\beta}}$ and the estimated variance-covariance matrix of $\widehat{\boldsymbol{\beta}}$, $\widehat{\mathrm{V}}_{\boldsymbol{\beta}}$, or to the original phenotypes and genotypes, $\underset{(N\times K)}{\mathrm{Y}}$ and $\underset{(N\times M)}{\mathrm{X}}$, and any other covariates that we want to regress out from the phenotypes.

The prior distribution for the null model, $p\left(\boldsymbol{\beta}|\mathrm{NULL}\right)$, is simply the point mass at $\boldsymbol{\beta} = 0$. In section 2, we show how we approximate the likelihood function for $\boldsymbol{\beta}$, $p\left(\mathrm{Data}|\boldsymbol{\beta}\right)$; in section 3, we define the prior distribution $p\left(\boldsymbol{\beta}|\mathrm{ALT}\right)$ for the alternative model; and finally, in section 4, we compute the BF.

#### 2 Likelihood function

A maximum likelihood estimator of B is given by the ordinary least-squares method

$$\widehat{\mathrm{B}} = \left(\mathrm{X}^\intercal\mathrm{X}\right)^{-1}\mathrm{X}^\intercal\mathrm{Y},$$

that in vectorized form is denoted by $\widehat{\boldsymbol{\beta}} = \left(\widehat{\boldsymbol{\beta}}_1, \ldots, \widehat{\boldsymbol{\beta}}_M\right)^\intercal$. An estimator of the variance-covariance of $\widehat{\boldsymbol{\beta}}$ is given by

$$\widehat{\mathrm{V}}_{\boldsymbol{\beta}} = \left(\mathrm{X}^\intercal\mathrm{X}\right)^{-1} \otimes \widehat{\mathrm{V}}_{\mathrm{Y}},$$

where $\widehat{\mathrm{V}}_{\mathrm{Y}}$ is the estimated residual variance-covariance matrix of Y given X.

Following Band et al. [1], we approximate the likelihood function of $\boldsymbol{\beta}$ by a multivariate normal distribution with mean $\widehat{\boldsymbol{\beta}}$ and variance-covariance matrix $\widehat{\mathrm{V}}_{\boldsymbol{\beta}}$. Note that by approximating $\widehat{\mathrm{V}}_Y$ via the trait correlation matrix, this likelihood approximation does not require access to the individual level data X and Y but only to the summary data of effect sizes $\widehat{\boldsymbol{\beta}}$, LD-matrix $\mathrm{X}^\intercal\mathrm{X}$, and a trait correlation estimate.

### 3 Prior of $\boldsymbol{\beta}$ in the alternative model

We construct the prior distribution $p(\boldsymbol{\beta}|\mathrm{ALT})$ for the alternative model in three steps, allowing the user to specify correlations between effects of different variants on different traits across different studies.

In a single study, the prior density for $\boldsymbol{\beta}$ incorporates the expected correlation of genetic effects among a group of variants ($\mathbf{R}_{\mathrm{var}}$) and among a group of phenotypes ($\mathbf{R}_{\mathrm{phen}}$). In addition, we incorporate an expected spread of the effect size of each variant by scaling $\mathbf{R}_{\mathrm{var}}$ as

$$\mathbf{S}_{\mathrm{var}} = \Delta(\sigma_m)\,\mathbf{R}_{\mathrm{var}}\Delta(\sigma_m)\,,$$

where $\Delta(\sigma_m)$ is a diagonal matrix with entries $\sigma_m$ determining the spread of the effect size distribution for each variant $m \leq M$. Thus, we can model settings where, e.g., protein-truncating variants have larger effect sizes ($\sigma = 0.5$) than missense variants ($\sigma = 0.2$). Note that when $\sigma_m = 1$ for all $m$, then $\mathbf{S}_{\mathrm{var}} = \mathbf{R}_{\mathrm{var}}$.

All in all, our prior density for $\boldsymbol{\beta}$ under alternative model is

$$\boldsymbol{\beta}|\mathrm{ALT} \sim \mathcal{N}(\mathbf{0}, \mathbf{U})\,, \text{ where } \mathbf{U} = \mathbf{S}_{\mathrm{var}} \otimes \mathbf{R}_{\mathrm{phen}}.$$

When we have data from multiple studies, we allow for possible differences in genetic effects across ethnicities or populations, extending the Approximate Bayes Factors of Band et al. [1] and the summary statistics approach of RAREMETAL [2] from univariate to multivariate phenotypes. Let $\widehat{\boldsymbol{\beta}} = \left(\widehat{\beta}_{s,m,k}\right) = \left(\widehat{\beta}_{1,1,1}, \widehat{\beta}_{1,1,2}, \ldots, \widehat{\beta}_{1,1,K}, \widehat{\beta}_{1,2,1}, \ldots, \widehat{\beta}_{1,2,K}, \ldots, \widehat{\beta}_{1,M,K}, \widehat{\beta}_{2,1,1}, \ldots, \widehat{\beta}_{S,M,K}\right)$, where $S$ is the number of studies, $M$ is the number of variants, and $K$ is the number of phenotypes. As with a single study, we incorporate the expected correlation of genetic effects between a pair of variants and a single phenotype using the matrix $\mathbf{S}_{\mathrm{var}}$, between a variant and a pair of phenotypes using the matrix $\mathbf{R}_{\mathrm{phen}}$, and we introduce the matrix $\mathbf{R}_{\mathrm{study}}$ to specify a prior on the similarity in effect sizes across the studies. Thus, the prior is

$$\boldsymbol{\beta} \sim \mathcal{N}(\mathbf{0}, \mathbf{U})\,, \text{ where } \mathbf{U} = \mathbf{R}_{\mathrm{study}} \otimes (\mathbf{S}_{\mathrm{var}} \otimes \mathbf{R}_{\mathrm{phen}})\,.$$

It is also straightforward to include a non-zero vector $\boldsymbol{\mu}$ as a prior mean of genetic effects, in which case the prior is

$$\boldsymbol{\beta} \sim \mathcal{N}(\boldsymbol{\mu}, \mathbf{U})\,.$$

We use this, for example, when screening for protective rare variants that have a pre-specified beneficial profile on a set of risk factors.

### 4 $\mathbf{BF_{MRP}}$

The Bayes Factor is the ratio of the marginal likelihoods between the alternative and the null model. The marginal likelihood for the alternative model is

$$\int_{\boldsymbol{\beta}} p(\mathrm{Data}|\boldsymbol{\beta})\, p(\boldsymbol{\beta}|\mathrm{ALT})\, d\boldsymbol{\beta} = c \times \mathcal{N}\left(\widehat{\boldsymbol{\beta}}; \boldsymbol{\mu}, \widehat{\mathrm{V}}_{\boldsymbol{\beta}} + \mathbf{U}\right)$$

and the marginal likelihood for the null model is

$$\int_{\boldsymbol{\beta}} p\left(\text{Data}|\boldsymbol{\beta}\right) p\left(\boldsymbol{\beta}|\text{NULL}\right) d\boldsymbol{\beta} = c \times \mathcal{N}\left(\widehat{\boldsymbol{\beta}}; 0, \widehat{\mathrm{V}}_{\boldsymbol{\beta}}\right).$$

The Bayes Factor is given by

$$\mathrm{BF}_{\mathrm{MRP}} = \frac{\det\left(\widehat{\mathrm{V}}_{\boldsymbol{\beta}} + \mathbf{U}\right)^{-\frac{1}{2}} \exp\left[-\frac{1}{2}\left(\widehat{\boldsymbol{\beta}} - \boldsymbol{\mu}\right)^{\mathsf{T}} \left(\widehat{\mathrm{V}}_{\boldsymbol{\beta}} + \mathbf{U}\right)^{-1} \left(\widehat{\boldsymbol{\beta}} - \boldsymbol{\mu}\right)\right]}{\det\left(\widehat{\mathrm{V}}_{\boldsymbol{\beta}}\right)^{-\frac{1}{2}} \exp\left[-\frac{1}{2}\widehat{\boldsymbol{\beta}}^{\mathsf{T}} \widehat{\mathrm{V}}_{\boldsymbol{\beta}}^{-1} \widehat{\boldsymbol{\beta}}\right]}.$$

When $\boldsymbol{\mu} = 0$, $\mathrm{BF}_{\mathrm{MRP}}$ is an increasing function of the following quadratic form

$$Q(\widehat{\boldsymbol{\beta}}; \widehat{\mathrm{V}}_{\boldsymbol{\beta}}, \mathbf{U}) = \widehat{\boldsymbol{\beta}}^{\mathsf{T}} \left(\widehat{\mathrm{V}}_{\boldsymbol{\beta}}^{-1} - (\widehat{\mathrm{V}}_{\boldsymbol{\beta}} + \mathbf{U})^{-1}\right) \widehat{\boldsymbol{\beta}}. \tag{1}$$

Furthermore, this quadratic form is the only part of the $\mathrm{BF}_{\mathrm{MRP}}$ that depends on $\widehat{\boldsymbol{\beta}}$. Thus, by deriving a distribution of $Q(\widehat{\boldsymbol{\beta}}; \widehat{\mathrm{V}}_{\boldsymbol{\beta}}, \mathbf{U})$ under the null model we can compute a p-value when $\mathrm{BF}_{\mathrm{MRP}}$ is used as a test statistic. According to basic properties of quadratic forms of Gaussian variables, $Q(\widehat{\boldsymbol{\beta}}; \widehat{\mathrm{V}}_{\boldsymbol{\beta}}, \mathbf{U}) \sim \sum_{i=1}^{n} d_i \chi_i^2$, where $\chi_i^2$ is an independent sample from a $\chi_1^2$ distribution (chi-square with one degree of freedom), and $d_i$ are the eigenvalues of matrix $I - (\widehat{\mathrm{V}}_{\boldsymbol{\beta}} + \mathbf{U})^{-1} \widehat{\mathrm{V}}_{\boldsymbol{\beta}}$. The distribution function for a mixture of chi-squares can be numerically evaluated by the R-package 'CompQuadForm' [3].

#### 4.1 MRP Bayes Factor computation

To compute the Bayes Factor

$$\mathrm{BF}_{\mathrm{MRP}} = \frac{\det\left(\widehat{\mathrm{V}}_{\boldsymbol{\beta}} + \mathbf{U}\right)^{-\frac{1}{2}} \exp\left[-\frac{1}{2}\left(\widehat{\boldsymbol{\beta}} - \boldsymbol{\mu}\right)^{\mathsf{T}} \left(\widehat{\mathrm{V}}_{\boldsymbol{\beta}} + \mathbf{U}\right)^{-1} \left(\widehat{\boldsymbol{\beta}} - \boldsymbol{\mu}\right)\right]}{\det\left(\widehat{\mathrm{V}}_{\boldsymbol{\beta}}\right)^{-\frac{1}{2}} \exp\left[-\frac{1}{2}\widehat{\boldsymbol{\beta}}^{\mathsf{T}} \widehat{\mathrm{V}}_{\boldsymbol{\beta}}^{-1} \widehat{\boldsymbol{\beta}}\right]},$$

we first consider the term inside the exponential function:

$$\mathcal{E}\left(\widehat{\boldsymbol{\beta}}, \boldsymbol{\mu}, \widehat{\mathrm{V}}_{\boldsymbol{\beta}}, \mathbf{U}\right) = \frac{1}{2}\widehat{\boldsymbol{\beta}}^{\mathsf{T}} \widehat{\mathrm{V}}_{\boldsymbol{\beta}}^{-1} \widehat{\boldsymbol{\beta}} - \frac{1}{2}\left(\widehat{\boldsymbol{\beta}} - \boldsymbol{\mu}\right)^{\mathsf{T}} \left(\widehat{\mathrm{V}}_{\boldsymbol{\beta}} + \mathbf{U}\right)^{-1} \left(\widehat{\boldsymbol{\beta}} - \boldsymbol{\mu}\right).$$

Since $\widehat{\mathrm{V}}_{\boldsymbol{\beta}}$ and $\mathbf{U}$ are typically defined through Kronecker products of smaller matrices, their inverses are easier to compute than the inverse of their sum. Hence we use Woodbury matrix identity to write

$$\mathcal{E}\left(\widehat{\boldsymbol{\beta}}, \boldsymbol{\mu}, \widehat{\mathrm{V}}_{\boldsymbol{\beta}}, \mathbf{U}\right) = \frac{1}{2}\widehat{\boldsymbol{\beta}}^{\mathsf{T}} \widehat{\mathrm{V}}_{\boldsymbol{\beta}}^{-1} \widehat{\boldsymbol{\beta}} - \frac{1}{2}\left(\widehat{\boldsymbol{\beta}} - \boldsymbol{\mu}\right)^{\mathsf{T}} \left(\widehat{\mathrm{V}}_{\boldsymbol{\beta}}^{-1} - \widehat{\mathrm{V}}_{\boldsymbol{\beta}}^{-1}\left(\mathbf{U}^{-1} + \widehat{\mathrm{V}}_{\boldsymbol{\beta}}^{-1}\right)^{-1} \widehat{\mathrm{V}}_{\boldsymbol{\beta}}^{-1}\right) \left(\widehat{\boldsymbol{\beta}} - \boldsymbol{\mu}\right).$$

To simplify the determinant calculation we write

$$\det\left(\widehat{\mathrm{V}}_{\boldsymbol{\beta}} + \mathbf{U}\right) = \det\left(\widehat{\mathrm{V}}_{\boldsymbol{\beta}}\right) \det\left(\mathbf{I} + \widehat{\mathrm{V}}_{\boldsymbol{\beta}}^{-1}\mathbf{U}\right).$$

The logarithm of the Bayes Factor is then

$$\log\left(\mathrm{BF}_{\mathrm{MRP}}\right) = -\frac{1}{2}\log\left(\det\left(\mathbf{I} + \widehat{\mathrm{V}}_{\boldsymbol{\beta}}^{-1}\mathbf{U}\right)\right) + \mathcal{E}\left(\widehat{\boldsymbol{\beta}}, \boldsymbol{\mu}, \widehat{\mathrm{V}}_{\boldsymbol{\beta}}, \mathbf{U}\right).$$

If studies do not share individuals, $\widehat{\mathrm{V}}_{\boldsymbol{\beta}}$ is a block-diagonal matrix

$$\widehat{\mathrm{V}}_{\boldsymbol{\beta}} = \left[\begin{array}{c|c|c|c} \widehat{\mathrm{V}}_{\boldsymbol{\beta}}^{1} & 0 & \cdots & 0 \\ \hline 0 & \widehat{\mathrm{V}}_{\boldsymbol{\beta}}^{2} & \cdots & 0 \\ \hline \vdots & & \ddots & \vdots \\ \hline 0 & 0 & \cdots & \widehat{\mathrm{V}}_{\boldsymbol{\beta}}^{S} \end{array}\right].$$

If studies share individuals, e.g., controls, we can take the approach of Cichonska et al. [4] to use summary level data to estimate the correlation structure of the non-diagonal blocks caused by overlapping individuals.
